# Assessing distribution changes of selected native and alien invasive plant species under changing climatic conditions in Nyeri County, Kenya

**DOI:** 10.1101/2020.08.25.265991

**Authors:** Julius Maina Waititu, Charles Ndegwa Mundia, Arthur W Sichangi

**Affiliations:** Institute of Geomatics, GIS and Remote Sensing, Dedan Kimathi University of Technology, Nyeri, Kenya

**Author notes:** Department of Spatial Planning, Kenyatta University, Nairobi, Kenya (Julius Maina Waititu). **Corresponding author** (JMW); (JMW).

## Abstract

Changes in climatic conditions increases the risks of native and alien taxa expanding in geographical range and causing habitat transformations. The role of climate change in enhancing bio-invasions in local natural environments need to be assessed to guide on effective species management policy formulations. In this present study, we used species presence records, predictor variables and an ensemble of General Circulation Models data to predict suitable ecological niches for five of the selected invasive plant species within Nyeri County, Kenya. We predicted species distributions under RCP2.6, RCP4.5, and RCP8.5 emission scenarios for the years 2050 and 2070. We analysed species distribution changes to identify invasive species requiring immediate management action. Our analysis indicated that three of the five study species were suitable in ~50% of the study area while the other two were suitable in ~30% under the current climate. *Lantana camara L.* and *Solanum campylacanthum Hochst. ex A. Rich* species would experience the largest range shift distance of ~6 – 10km and the largest habitat gain of ~12 – 33% in the future. *Caesalpinia decapetala (Roth) Alston, Opuntia stricta (Haw.) Haw.* and *Senna didymobotrya (Fresen.) H.S. Irwin & Barneby* species on the other hand would have a decline in habitat range under future climate change scenarios. Although, *S. didymobotrya* is considered a native species, it would lose half of its current suitable habitat in the future. Range shift analysis showed all study species would generally shift to the north west direction or towards the Aberdare ranges. From this study we conclude that **i**nvasive species management programs for smaller geographical areas ought to consider projecting species distributions under climate change scenarios to identify areas with high possible biodiversity changes. This would be important to conservationists when prioritizing management actions of invasive species in the region where data on invasive species is still limited.

## Introduction

According to Richardson et al. [1], Naturalized plants are alien plants that have consistently reproduced with no human intervention over many growth periods while Invasive plants are naturalized plants that are able to produce many reproductive offsprings at considerable distances from parent plants. Many of the alien invasive plant species have profound negative impacts on forest resources, water resources, and agricultural ecosystems [2–4]. In some instances, alien invasive species may be beneficial to new environments e.g. supporting local fauna in their habitats, reducing carbon footprint, and provision of firewood [5].

Native species are considered a nuisance if they are expanding in range and causing habitat transformations [6]. Although proliferation of alien species in new environments may be attributed to lack of natural enemies inhibiting their survival [7], changes in climatic conditions may render any of the alien taxa to extinction or may enable its spread and survival [1]. To the advantage of alien invasive species, their ability to adapt in habitats with varying climatic conditions has contributed to their expansion in geographical range [8].

Conservationists need to project distributions of invasive species during the temperature overshoot periods i.e. temperatures over 1.5°C in the course of 21^st^ century so as to determine species survival boundary limits to guide management strategies. In the Intergovernmental Panel on Climate Change (IPCC) 5^th^ Assessment Report [9], time-dependent projections of atmospheric greenhouse gas (GHG) concentrations are described in three Representative Concentration Pathways (RCPs) namely; the stringent mitigation scenario (RCP2.6) denoting a peak and decline of temperatures below 1.6°C, the intermediate stages (RCP4.5 and RCP6.0) denoting a stabilization without overshoot, and the higher GHG emission scenario (RCP8.5) denoting rising temperatures. As described in IPCC [9 p60] and [10], the period 2016 – 2035 for all RCP scenarios is predicted in the range of 0.3°C −0.7°C similar to 1986 – 2005 reference period. The mean temperatures under RCP2.6, RCP4.5 and RCP8.5 for the period 2046-2065 are predicted to be 1.0°C, 1.4°C, and 2.0°C respectively while for the period 2081 – 2100, mean temperatures will be 1.0°C, 1.8°C and 3.7°C respectively. Warming temperature of 2°C and above is expected to exacerbate risks brought about by spread of invasive plant species than that maintained at 1.5°C and below [11].

Management decisions on invasive species usually depend on their impacts on native vegetation diversity and richness [12]. Although it may be challenging to quantify such impacts if species prevalence is unknown [13], building Species Distribution Models (SDM) forms an integral part in estimating prevalence and distribution changes. SDMs have been built on various biogeographical scales using correlative techniques relying on species occurrence data and predictor variables [14–16]. SDMs carried out at large biogeographical scales e.g. at, regional, national or continental level provide baseline information for conservation management policy formulation [17,18] while those at a local scale are used to provide estimates on community level species population for immediate management prioritization [19].

Witt et al. [6] spurred interests in documenting invasive species through a broad baseline study on multiple invasive species status in East Africa region. Some of their study species included: *Lantana camara L., Opuntia Stricta (Haw.) Haw*, and *Acacia mearnsii De Wild* which are listed in the land plant category of 100 worst invasive alien species [20]. *L. camara* prevalence in most parts of East Africa has led to biodiversity and livelihood losses [18]. *O. Stricta* on the other hand is naturalized and invasive in arid and semi-arid areas of Africa. In Laikipia county, Kenya, 17% of the area is invaded [21]. According to Witt et al. [6], only south Africa has a comprehensive up-to-date database on invasive species. Gichua et al. [22] noted that information on the pattern of introduction and the spread dynamics of invasive species in Kenya is insufficient for effective monitoring and management of invasive species. In highland protected areas of Rwanda, *A. mearnsii* has survived as an understory in pine and eucalyptus plantations and has generally done well in altitudes above 1200m including those outside natural forests [23]. *Caesalpinia decapetala (Roth) Alston* has been studied extensively in South Africa where it has had invasive presence in natural areas such as forests and riparian reserves as well as in grazing areas where their prickles injure livestock [24]. In East Africa, distribution maps of *L. camara* from the work of Witt et al. [6] shows invasive status in central and western parts of Kenya and that of *Solanum campylacanthum Hochst. ex A. Rich.* is widespread along roadsides, disturbed areas and forest edges [25].

Review of current literature indicate limited current and future species distribution studies particularly at a local scale despite their importance in conservation management strategies. For instance, in Nyeri County where two of the most important biodiversity-rich ecosystems i.e. the Aberdare national park and the Mt. Kenya national park and forest ecosystem [26] are located, future species distribution studies have not been conducted. Uncontrolled spread of invasive species leads to degradation of natural habitats. Such impacts have been reported in Mt. Marsabit forest where ecosystem degradation has resulted to loss of forest cover from initial 18,363 hectares in 1973 to 11,000 hectares by 2013 [27].

This study, aimed at providing a species distribution modelling framework to estimate the current and future species distribution at a local scale level to support invasive species policy formulation. The objectives of this study were broken into two: (i) To demonstrate the applicability of selected General Circulation Models (GCM) data for the year 2050 and 2070 under RCP2.6, RCP4.5, and RCP8.5 in ensemble species distribution modelling; and (ii) To assess habitat suitability changes of the study species under current and future climate change scenarios.

## Materials and methods

### Study area and its ecological importance

This study was carried out in Nyeri County in central Kenya covering an area of ~ 3278.16 Km^2^. It is strategically located between Mt. Kenya ecosystem to the east and Aberdare ecosystem to the west within Latitudes 0° 38’ 45” S and 0° 0’ 42” S and Longitudes 36° 35’ 28” E and 37° 18’ 29” E (Fig 1). These two ecosystems and other isolated forest hills play a vital role in the climate of the area and serve as wildlife habitats, forest reserves and water catchment areas [28]. Nyeri County is comprised of Kieni, Othaya, Mathira, Mukurwe-ini, Tetu and Nyeri Town sub-counties (Fig 1). It has four agro climatic zones namely: Humid (I), Sub humid (II), Semi humid (III), Semi humid to Semi-arid (IV) and Semi-Arid (V). Kieni sub-county falls in II, III, IV & V zones, Othaya, Nyeri Town and Tetu sub-counties fall in zones I & II, while Mathira and Mukurwe-ini sub-counties fall in zones I, II and III [29]. On average, annual rainfall in Nyeri ranges between 1200 – 1600mm and 500 – 1500mm during long and short rains respectively while the monthly mean temperatures ranges between 12 – 21°C [28].

**Fig 1.**
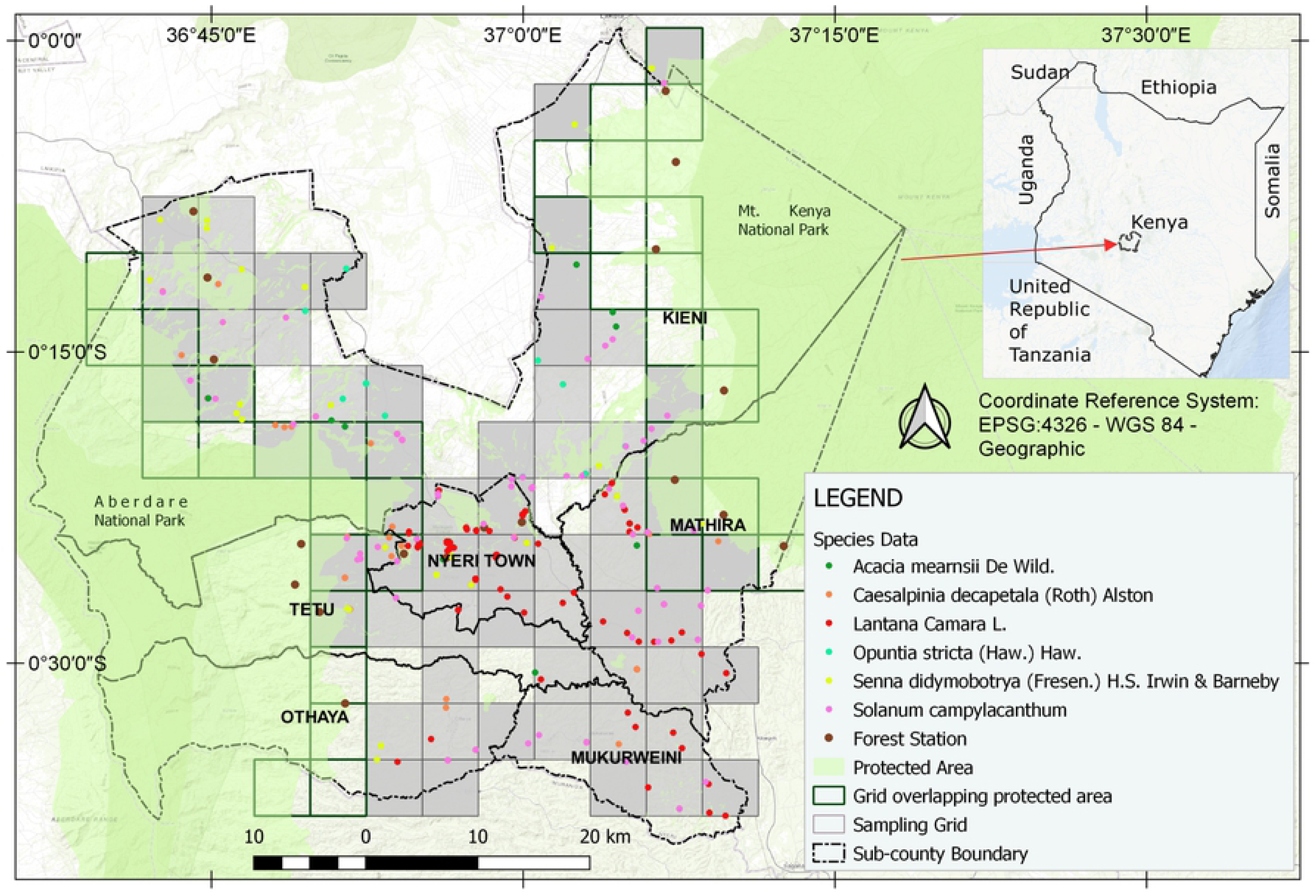
Study area map showing distribution of invasive species field occurrence records.

Invasive species such as C. decapelata, S. campylacanthum, L. camara L, Datura dothistroma, Resinus communis, Fraxinus pennysilvania, A. mearnsii and Rubus steudneri have invaded Mt. Kenya and the Aberdare forest ecosystems [30]. Among these species, A. Mearnsii, C. decapetala (Roth) Alston, C. decapetala, S. campylacanthum have been reported by Witt et al. [6] to be among invasive species expanding in range and having the greatest impacts on habitats such as forests and arable land. Management of spread of invasive species is therefore required in Nyeri to mitigate the threats posed by invasive species on biodiversity within natural habitats. Arable land need to be protected from invasive species since agriculture is the main economic activity in Nyeri county [28].

### Collection of species occurrence records

Field survey records for six invasive species namely: *L. camara, C. decapetala, Senna didymobotrya (Fresen.) H.S. Irwin & Barneby, S. campylacanthum, O. stricta* and *A. mearnsii* were collected along selected road networks in between October 2019 and February 2020 using a handheld GPS receiver (±3 m accuracy).

Since our work involved estimation of target species population, sampling species records through roadside surveys provided adequate estimation at a cheaper cost [31]. Furthermore, transport routes have been associated with introduction and eventual spread of invasive species [32,33] and therefore act as a factor influencing dispersal limitation [19]. Our target species also invades roadside habitats [25] and therefore probability of sighting them along survey routes was considered very high. Since forest edges provide favourable biotic and abiotic conditions for alien invasive species introduction and eventual spread to fragmented forest interiors [34], routes leading to forest edges were also considered.

As we drove along survey routes, target species occurrence locations were collected at approximately 2 to 5 km successive intervals. We selected a maximum of 5 km sampling interval to increase sampling intensity. The choice of interval was subjective as there are no reference standard intervals. For instance, [35] and [36] used 5-10km while [37] considered 25km interval indicating the subjective nature on selection of intervals. A shorter interval of 2km was considered on routes with steeper terrains to account for vegetation diversity changes as elevation increases [35]. At each stop, species occurrence locations were collected on both sides of the road and in the adjacent habitats if any of the target species was sighted.

In addition, a geographical grid of 5 × 5 Km^2^ over the entire study area was generated. The grid cell size was chosen to align with the maximum roadside survey interval chosen for this study. The grid cells served the purpose of estimating prevalence of target species by estimating the percentage distribution of present grid cells to the total surveyed grid cells [38] (Table 1 in S1 Appendix, Fig 1 and Table 1). Sampled grid cells depicted in grey colour in Fig 1 were all the sampled cells containing at least 1 of the species occurrence. In addition, the use of these grid cells served as background extent in generating background data with the same bias as the collected presence records [39].

**Table 1.** Description of study species, their prevalence and number of records used in SDM. Species scientific, family and common name, life form, and origin were adopted from Witt & Luke [25].

| Species and Family name | Common name | Life form | Origin | Class | Species presence records | Presence cells | Sampled grid cells out of 109 sampled cells | Species prevalence | Minimum records for SDM | Rarefied records |
| --- | --- | --- | --- | --- | --- | --- | --- | --- | --- | --- |
| *Acacia mearnsii* De Wild. (Fabaceae) | black wattle | Tree or shrub | South-eastern Australia and Tasmania. | Alien | 11 | 8 | 7% | 0.07 | 8 | 9 |
| *Caesalpinia decapetala* (Roth) Alston (Fabaceae) | Mauritius or Mysore thorn | Evergreen shrub / Climber | Native of Asia (India, Sri Lanka, China, Japan & Malaysia). | Alien | 37 | 22 | 20% | 0.20 | 19 | 26 |
| *Lantana camara* L. (Verbenaceae) | Lantana, tickberry | Tree or Shrub | Subtropical and tropical America. | Alien | 141 | 30 | 33% | 0.33 | 25 | 60 |
| *Opuntia stricta* (Haw.) Haw. (Cactaceae) | Erect prickly pear | Succulent tree or shrub | South-east USA, eastern Mexico and some Caribbean Islands. | Alien | 17 | 12 | 11% | 0.11 | 14 | 14 |
| *Senna didymobotrya* (Fresen.) H.S. Irwin & Barneby (Fabaceae) | African senna | Tree or shrub | Tropical Africa | Native | 43 | 27 | 24% | 0.24 | 19 | 31 |
| *Solanum campylacanthum* Hochst. ex A.Rich. (Solanaceae) | Bitter apple, Sodom apple | Shrub | Africa, Middle East and India. | Native | 65 | 34 | 31% | 0.31 | 25 | 50 |

### Modelling predictor variables

We considered a set of predictor variables accounting for abiotic and biotic factors (see Table 2 in S1 Appendix.). The 19 standard WorldClim bioclimatic variables derived from averaged climate data for the period 1970 – 2000 [40] at a spatial resolution of 30 arc seconds (~1 km^2^) were downloaded from WorldClim (https://worldclim.org/). Global aridity index and global potential evapo-transpiration predictor variable layers constituting water balance variables were obtained from Trabucco and Zomer [41]. Digital Elevation Model (DEM) data at a spatial resolution of 12.5 × 12.5m was downloaded from Alaska DEM Facility [42]. Slope, aspect, plan and profile curvatures and topographic wetness index (twi) were derived from the Alaska DEM data. Landcover data derived from sentinel 2 imagery data was downloaded from ESA (2017) while Normalized difference moisture index (NDMI) and Normalized difference vegetation index (NDVI) were generated from January 2019 sentinel 2 imagery data downloaded from Copernicus Open Access Hub (https://scihub.copernicus.eu/dhus). Kenya Soil data (soil pH and soil class) was obtained from Hengl et al. [44] in raster format. All predictor variable layers were resampled to 30 arc seconds spatial resolution and masked to study area extents (Fig 1).

**Table 2.** Retained noncollinear predictor variables (r < 0.70) for individual study species.

| *C. decapetala* | *L. camara* | *O. stricta* | *S. didymobotrya* | *S. campylacanthum* |
| --- | --- | --- | --- | --- |
| **Aspect<sup>a</sup>** | Aspect | **Aridity index<sup>a</sup>** | Aspect | Aspect |
| bio14 (Precipitation of Driest Month) | bio14 (Precipitation of Driest Month) | **Aspect<sup>a</sup>** | **bio18 (Precipitation of Warmest Quarter)<sup>a</sup>** | bio14 (Precipitation of Driest Month) |
| bio15 (Precipitation Seasonality (Coefficient of Variation)) | bio18 (Precipitation of Warmest Quarter) | bio1 (Annual Mean Temperature) | bio4 (Temperature Seasonality) | **bio18 (Precipitation of Warmest Quarter)<sup>a</sup>** |
| **bio18 (Precipitation of Warmest Quarter)<sup>a</sup>** | **bio4 (Temperature Seasonality)<sup>a</sup> (SD × 100)** | **bio14 (Precipitation of Driest Month)<sup>a</sup>** | **bio6 (Min Temperature of Coldest Month)<sup>a</sup>** | **bio3 (Isothermality)<sup>a</sup>** |
| bio2 (Mean Diurnal Range) | bio7 (Temperature Annual Range) | bio18 (Precipitation of Warmest Quarter) | **bio7 (Temperature Annual Range)<sup>a</sup>** | bio7 (Temperature Annual Range) |
| bio3 (Isothermality) | **DEM<sup>a</sup>** | **bio4 (Temperature Seasonality)<sup>a</sup> (SD × 100)** | landcover | **bio9 (Mean Temperature of Driest Quarter)<sup>a</sup>** |
| **bio6 (Min Temperature of Coldest Month)<sup>a</sup>** | landcover | bio6 (Min Temperature of Coldest Month) | **NDMI<sup>a</sup>** | **landcover<sup>a</sup>** |
| **landcover<sup>a</sup>** | NDMI | landcover | NDVI | NDMI |
| NDMI | NDVI | NDMI | Plan curvature | **NDVI<sup>a</sup>** |
| NDVI | Plan curvature | NDVI | Profile curvature | Plan curvature |
| Plan curvature | Profile curvature | Plan curvature | slope | Profile curvature |
| Profile curvature | slope | **Soil Class<sup>a</sup>** | Soil Class | **slope<sup>a</sup>** |
| **slope** | Soil Class |  | twi | Soil Class |
| Soil Class | Soil pH |  |  | Soil pH |
| **Soil pH<sup>a</sup>** | twi |  |  | twi |
| **twi<sup>a</sup>** |  |  |  |  |

twi, topographic wetness index; NDMI, Normalized Difference Moisture Index; NDVI, Normalized Difference Vegetation Index; DEM, Digital Elevation Model.
<sup>a</sup>Predictor variables with relative variable importance (correlation test: ' $1 - correlation$ ' > 0.05) for both maxent and random forest algorithms.

Spatial downscaling future climate data representing future emission pathways for the year 2050 (2040-2069) and 2070 (2060–2089) [45] at 30 arc seconds spatial resolution was downloaded from (http://www.ccafs-climate.org/). We considered 6 Coupled Model Intercomparison Project Phase 5 (CMIP5) GCM data namely: BCC-CSM1.1(m), GFDL-ESM2G, HadGEM2-ES, IPSL-CM5A-MR, MIROC-ESM-CHEM, and NCAR-CCSM4 under the RCP2.6, RCP4.5 and RCP8.5. According to McSweeney, Jones, Lee, and Rowell [46], these GCM models performance ratings on replicating timings for annual precipitation and temperature cycles are relatively the same with no significance differences in the Horn of Africa region within which our study area lies. Researchers in ecological niche modelling have recommended use of an ensemble of values averaged for selected GCMs e.g. [47,48] to account for differences in climate predictions of individual GCMs. We used individual GCM data to predict future habitat suitability for study species and thereafter assessed spatial similarities between individual binary map outputs to determine candidate GCM models for an ensemble distribution modelling.

### Predictor variables multi-collinearity analysis

A total of 32 model predictor variables (Table 2 in S1 Appendix) were subjected to multi-collinearity test using pearson’s correlation coefficient (r) and variance inflation factor (VIF). Calculation of VIF was as shown in Equation (1). Assessing and removing collinear variables conforms with statistical assumptions in regression models [49].

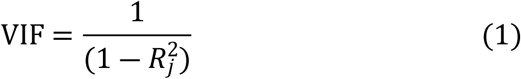

where 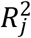 is the coefficient of determination derived from model variables *j* [50].

We implemented this in R statistical software [51] and the ‘*usdm’* package [52]. We used the presence records for individual species to extract predictor values from all predictor variables and converted them into a matrix data frame for multi-collinearity test. Predictor variables with VIF > 10 [53] and r > 0.70 [13] were excluded. Retained predictor variables were as shown in Table 2.

### Preparation of presence and background data

#### Rarefying presence samples

We used the SDMtoolbox 2.4 for ArcGIS 10.5 [54] to spatially rarefy individual species occurrence data. Spatially clustered points introduces environmental biases and tend to affect model’s ability to predict given new data [54]. We used 1 km Euclidean distance so as to match the predictor variables 1 Km^2^ spatial resolution [55]. A total of 190 records out of 314 were retained for model fitting (Table 1).

#### Minimum sample size for generating accurate SDM

Filtered species presence data (Table 1) were assessed to determine whether the minimum number was met for building accurate SDMs [13]. Five of the study species except for *A. mearnsii* were found to be within the recommended prevalence range (0.1 (10%) to 0.5 (50%)) as shown in Table 1. R scripts adopted from van Proosdij et al. [13] were run and results based on real AUC threshold (AUC > 0.9) from the maxent model built using linear and quadratic features were considered to assess the required minimum sample records. The minimum records were 8, 11, 15, 19, 19 and 25 for 0.05, 0.10, 0.20, 0.30, 0.40, and 0.50 prevalence values respectively (Table 3 in S1 Appendix). *A. mearnsii* had 9 presence records and a prevalence of < 0.10 and was therefore dropped from further distribution modelling as recommended by van Proosdij et al. [13].

**Table 3.** Average AUC, TSS, maxSSS model accuracy assessment values based on 30% subsampled dependent test data and Null model AUC values used to assess significance of species model AUC values.

| Species<br>scientific<br>name | Species model AUC |  |  | Null-model<br>AUC | Mean TSS |  |  | Optimized mean threshold (TR) using<br>(maxSSS) criteria |  |  |
| --- | --- | --- | --- | --- | --- | --- | --- | --- | --- | --- |
|  | rf | maxent | Mean |  | rf | maxent | Mean | rf | maxent | Mean |
| *C. decapetala* | 0.75 | 0.77 | 0.76 | 0.71 | 0.56 | 0.63 | 0.59 | 0.49 | 0.44 | 0.46 |
| *L. camara* | 0.90 | 0.91 | 0.91 | 0.66 | 0.79 | 0.78 | 0.79 | 0.47 | 0.39 | 0.43 |
| *O. stricta* | 0.94 | 0.92 | 0.93 | 0.71 | 0.87 | 0.83 | 0.85 | 0.49 | 0.49 | 0.49 |
| *S. didymobotrya* | 0.76 | 0.76 | 0.76 | 0.69 | 0.58 | 0.58 | 0.58 | 0.30 | 0.42 | 0.36 |
| *S. campylacanthum.* | 0.75 | 0.79 | 0.77 | 0.66 | 0.53 | 0.57 | 0.55 | 0.44 | 0.40 | 0.42 |

rf, random forest;

#### Generating background data

Background data were generated randomly within study area sampling grid cells (Fig 1). Based on the number of rarefied presence data (see Table 1, S6 Dataset), a ratio of 1:1 for presence/background data for *C. decapetala (Roth) Alston, L. camara* and *S. campylacanthum* species and a ratio of 2:1 to presence data for *O. stricta* and *S. didymobotrya* species were used to fit models with random forest and maxent methods. According to Barbet-Massin et al. [56], the same number of background data selection should be used for machine learning methods. However, the effect of using large background data was reported in the work of Liu, Newell, and White [57] where low model accuracies were found for random forest models built with large number of background samples (n=5000). They used a ratio of 2:1 to presence data to improve on species model AUC values.

### Current and future species distribution modelling

#### Model fitting and evaluation

As illustrated in Fig 2 and previous sections, SDMs were generated and evaluated using R statistical software [51] and the ‘*sdm*’ package [53]. SDM models were built with maxent [58], and Random Forest [59] methods using presence/background data and noncollinear predictor variables (Table 2). Random forest algorithm has gained popularity in ecological niche modelling due to its good performance as reported in Zhang et al. [60] and Shabani, Kumar, and Ahmadi [61] while the Maxent algorithm is popular due to its high predictive power than the traditional logistic regression [14] regardless of available sample size [17]. Species presence data were randomly split into two sub-sets; 70% for model fitting and 30% for testing community level predictions. Using all datasets may cause model overfitting and consequently fail to provide generality of the model for future time predictions [62]. We used 10 runs of subsample replications for both maxent and random forest algorithms hence generating 10 models for both. We used linear and quadratic features, a regularization multiplier of 1, with extrapolation and clamping to fit maxent models and retained defaults for R package ‘randomForest’ [63] models. A regularization multiplier of 1 provided a balance between a largely spread out and a localized prediction. Clamping helped in maintaining the range of predictor values seen during model training when predicting distributions with novel climate data [64].

**Fig 2.**
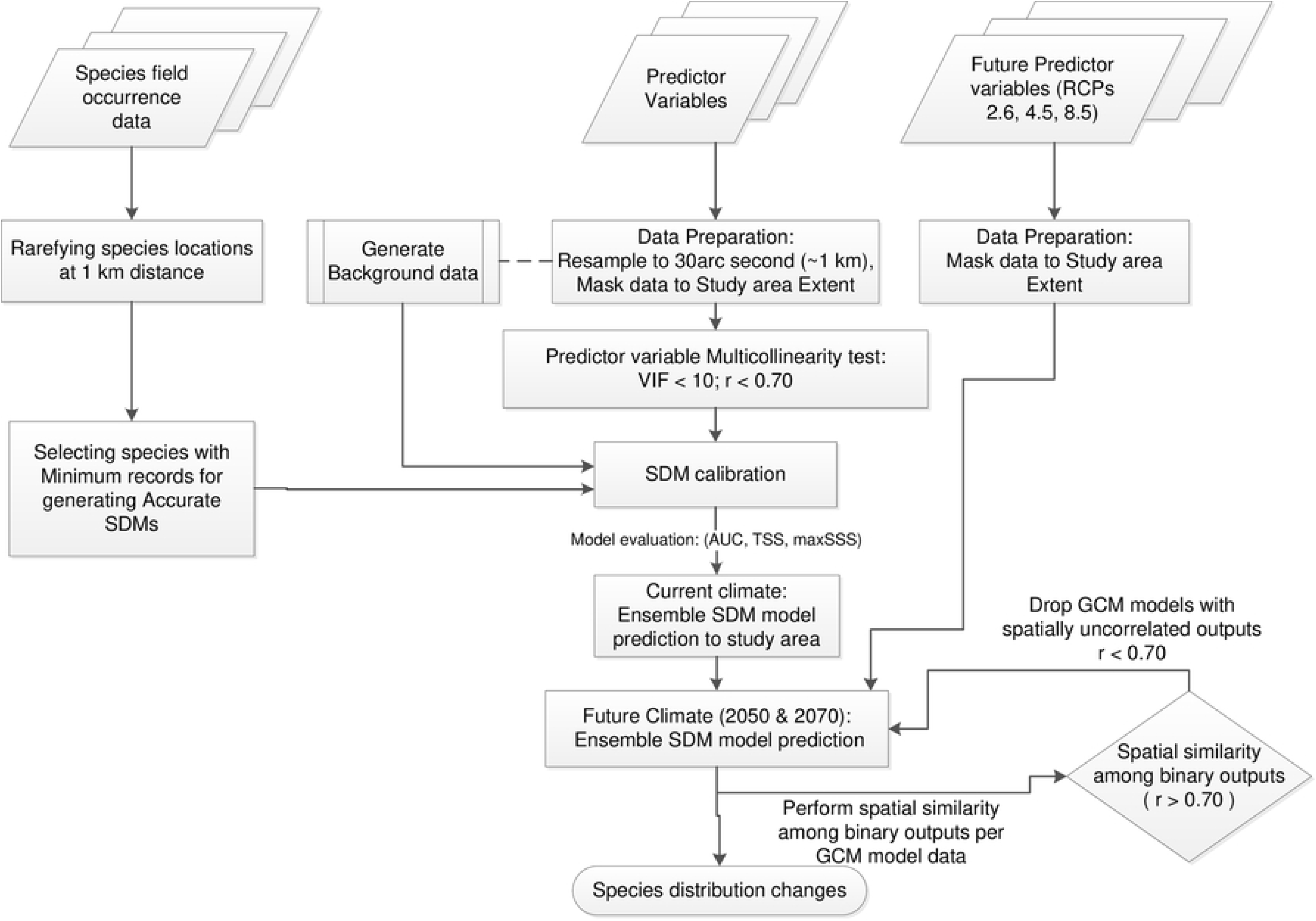
A conceptual framework for species distribution modelling under current and future climate scenarios.

**Table 4.** Percentage ensemble distribution changes for current climate period (1970-2000) and future climate periods; years 2050 (2040-2069) and 2070 (2060 - 2089).

| species<br>scientific<br>name | Year | range expansion (%) |  |  | no occupancy (absence<br>in current and future)<br>(%) |  |  | no change (present in<br>in current and future)<br>(%) |  |  | range contraction (%) |  |  | Range shift (km)<br>(%) |  |  | Shift<br>direct<br>ion<br>(%) |
| --- | --- | --- | --- | --- | --- | --- | --- | --- | --- | --- | --- | --- | --- | --- | --- | --- | --- |
|  |  | *a* | *b* | *c* | *a* | *b* | *c* | *a* | *b* | *c* | *a* | *b* | *c* | *a* | *b* | *c* |  |
| *C.*<br>*decapetala* | 2050 | 1.15 | 1.15 | 0.26 | 49.28 | 49.28 | 50.17 | 42.92 | 43.47 | 39.44 | 6.65 | 6.10 | 10.13 | 1.31 | 1.21 | 0.76 | NW |
|  | 2070 | 1.15 | 0.99 | 1.02 | 49.28 | 49.44 | 49.41 | 43.08 | 42.42 | 42.63 | 6.49 | 7.14 | 6.94 | 1.31 | 1.31 | 1.08 | NW |
| *L. camara* | 2050 | **12.14** | **12.77** | **14.87** | 57.21 | 56.58 | 54.49 | 30.59 | 30.59 | 30.62 | 0.05 | 0.05 | 0.03 | **8.42** | **8.91** | **9.65** | NW |
|  | 2070 | **11.62** | **12.35** | **11.54** | 57.73 | 57.00 | 57.81 | 30.62 | 30.59 | 30.57 | 0.03 | 0.05 | 0.08 | **7.99** | **8.49** | **8.08** | NW |
| *O. stricta* | 2050 | 0.52 | 0.92 | 0.31 | 69.41 | 69.01 | 69.62 | 27.06 | 27.87 | 19.47 | 3.01 | 2.20 | **10.60** | 1.49 | 1.37 | **5.64** | NW |
|  | 2070 | 1.20 | 0.60 | 0.71 | 68.73 | 69.33 | 69.22 | 28.42 | 27.03 | 27.51 | 1.65 | 3.04 | 2.56 | 1.32 | 1.30 | 1.12 | NW |
| *S.*<br>*didymobotry*<br>*a* | 2050 | 1.94 | 1.94 | 2.30 | 45.28 | 45.28 | 44.91 | 26.54 | 29.34 | 32.40 | **26.25** | **23.45** | **20.39** | 4.41 | 3.55 | 3.40 | NW |
|  | 2070 | 2.22 | 1.94 | 1.86 | 44.99 | 45.28 | 45.35 | 26.75 | 26.80 | 25.39 | **26.04** | **25.99** | **27.40** | 3.89 | 4.45 | 4.45 | W |
| *S.*<br>*campylacant*<br>*hum.* | 2050 | **33.16** | **33.11** | **25.52** | 19.03 | 19.08 | 26.67 | 47.61 | 47.53 | 46.82 | 0.21 | 0.29 | 0.99 | **6.69** | **6.59** | **5.52** | NW |
|  | 2070 | **33.84** | **32.27** | **30.25** | 18.35 | 19.92 | 21.93 | 47.74 | 47.66 | 47.76 | 0.08 | 0.16 | 0.05 | **6.69** | **6.59** | **6.52** | NW |

*a*, RCP2.6; *b*, RCP4.5; *c*, RCP8.5; NW, North West); W, West;
**Bold Values** indicate species range expansion or range contraction of $\geq 10\%$ and range shift of $\geq 5$ km).

Area under the receiver operating characteristics (ROC) curve (AUC) [38,65] and the True Skill Statistic (TSS) [66] average model assessment metrics were used for model evaluation (Table 4 in S1 Appendix). Average AUC ≥ 0.70 and TSS ≥ 0.5 from our fitted models were considered satisfactory for ensemble prediction to the rest of the study area. Average maximum Sensitivity plus Specificity (maxSSS) threshold values [67] were used to transform current and future probabilistic distribution maps into binary outputs.

We computed the species null-model AUC values to test significance of each of the obtained species model AUC values from random expectation [68]. A modification of the NullRandom function in ‘*dismo*’ package [69] was used to generate null-model AUC values for our study species. The randomly drawn collection localities (presences) within our geographical extent were the same number as the actual number of individual species records used in building our SDM models. We limited the number of background points to not more than 10,000 and repeated the randomizations 99 times. Comparison between null-model and species model AUC values showed all model AUC values were significantly better than those obtained by random chance (Table 3).

#### Ensemble predictions

Ensemble model predictions over entire study area using current climate data (Table 2) and future GCM climate data under climate change scenarios for the years 2050 and 2070 were based on, a weighted average over all fitted models for both maxent and random forest methods using the TSS statistic and maxSSS threshold criteria in the ‘*sdm’* R package [53].GCM outputs spatial similarity analysis

We used r and p-values in the function “*rcorr*” in “*Hmisc*” R package [70] to assess spatial similarity levels among pairs of individual GCM binary outputs per study species [71]. Spatial correlation analysis was based on a threshold of r ≥ 0.70 to indicate a ‘strong’ spatial correlation between pairs of GCM binary outputs per study species. Future predicted maps for the 3 RCPs and individual 6 GCMs for both future periods were taken as the variables (Table S4.5). Strongly correlated GCM model binary output maps for a given study species were selected as candidates for averaging into an ensemble of future predictor variables.

#### Change distribution analysis

Change distribution analysis between current and future binary output maps were performed in SDMtoolbox v2.4 [54]. We obtained change distributions in Km^2^ grouped into four classes namely: range expansion, no occupancy (absence) in both current and future, no change (presence) in current and future, and range contraction. To maintain consistency in the change distribution analysis, all SDM binary rasters were projected to an equal area projected coordinate system (Lambert Azimuthal Equal Area for Equatorial Region). Centroids for the current and future binary output maps were derived for calculation of species range shift distance in kilometres and direction shifts in bearings.

## Results

### Model performance

Results for the relative predictor variable importance and model performance are summarized in Table 2 and Table 3 respectively. All maxent and random forest models showed high AUC values ranging between 0.76 – 0.93 and were consequently significant above the null-model AUC values.

Relative variable importance as applied in determining species habitat suitability differed among study species (Fig 1 in S2 Appendix). Important bioclimatic variables among study species were generally <3 in addition to different topographic variables for each species. Precipitation of the warmest quarter(bio18) variable was common to *C. decapetala (Roth) Alston, S. didymobotrya*, and *S. campylacanthum* while the variable on temperature seasonality (bio4) was common to *L. camara* and *O. stricta.* Analysis of species response curves (Fig 2 in S2 Appendix) showed values ranging from ~250 – 350mm for bio18 enhanced *C. decapetala* distribution indicating that areas with precipitation of below 250mm during warmest quarter reduces its distribution. *S. didymobotrya* distribution declined with increasing bio18 values ranging between ~250 −375mm meaning that the species distribution does not depend on precipitation above this range while *S. campylacathum* responded positively within precipitation range of ~250 – 375mm during the warmest quarter. Probability of *L. camara* species distribution showed an increase within 6.5 – 11 bio4 values giving an indication that it survives well within a wide range of temperature variations throughout the year while values of ~6.0-7.0 increased *O. stricta* species response and values above 7.0 gradually decreased its response.

### Current habitat suitability maps output

Individual species binary distribution maps (Fig 3) showed that 50%, 31%, 30%, 53%, and 48% of the total geographical area as suitable for *C. decapetala* (Fig 3a)*, L. camara* (Fig 3b)*, O. stricta* (Fig 3c)*, S. didymobotrya* (Fig 3c) and *S. campylacanthum* (Fig 3d) species respectively. *O. stricta*species current suitable areas occupied most of Nyeri Town, parts of mukurwe-ini sub-county and in the semi-humid to semi-arid area of Kieni sub-counties. *Senna* species recorded the highest suitability area within Kieni sub-county and the least in Mukurwe-ini sub-county. *L. camara* and *O. stricta*species had relatively the same current suitable area with *L. camara* species covering the whole of Mukurwe-ini and Nyeri Town and parts of Othaya, Tetu, Mathira and Kieni sub-counties. *C. decapetala* and *S. campylacanthum* species suitable area covered half of the study area and were generally spread across all sub-counties. Notably, part of the suitable areas for *C. decapetala* species covered parts of the Aberdare ecosystem to the west. Other than *C. decapetala*, the rest of the species current suitable habitats were not predicted in the Mt. Kenya national reserve and the Aberdare national reserve.

**Fig 3.**
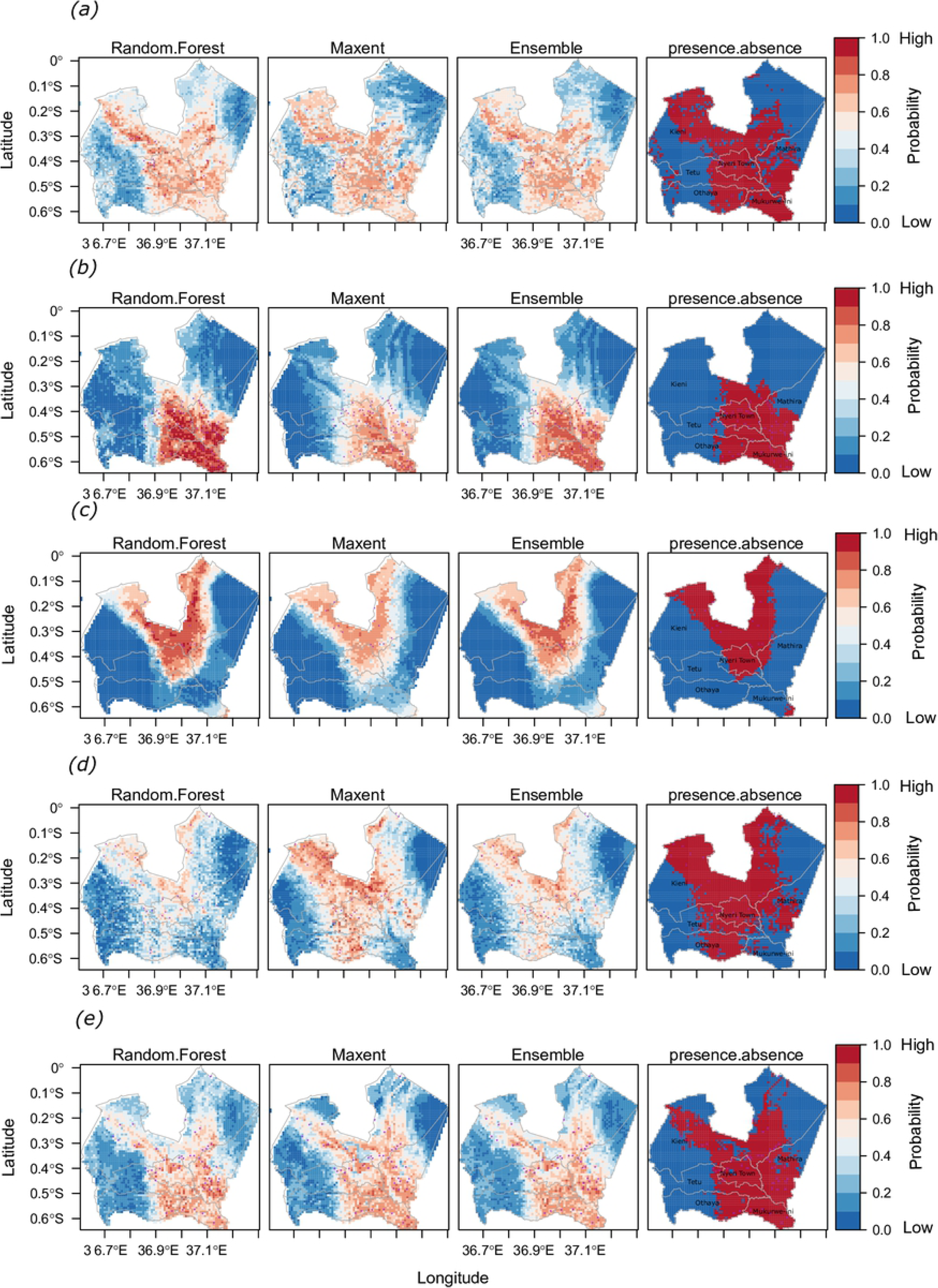
Species current distribution maps obtained with random forest and maxent algorithms and ensemble binary maps obtained in a weighted average. (a) C. decapetala (Roth) Alston; (b) L. camara; (c) O. stricta (Haw.) Haw.; (d) S. didymobotrya; (e) S. campylacanthum

### Future habitat suitability outputs

#### Spatial correlation among change distribution maps

Correlation coefficient values among binary output pairs of respective GCMs under the 3 RCPs for years 2050 and 2070 were found to be significantly different than 0 (*p < 0.001*). Spatial correlations among predicted binary outputs from all the six GCM (Table 1 in S5 Appendix) for *C. decapetala*, *L. camara* and *O. stricta*were above the *r* threshold value (*r* = (0.82-0.96); *r* = (0.86-0.98); and *r* = (0.71-0.97) respectively) (Fig 1 in S3 Appendix, Fig 1 in S4 Appendix).

Output binary maps for *S. didymobotrya* species were strongly correlated except for three binary output maps. These were correlations between BCC-CSM1.1_m and MIROC-ESM-CHEM outputs under RCP4.5 for year 2050 (*r* = 0.66) and for year 2070 (*r* = 0.62) and the lowest correlation value (*r* = 0.58) given by correlations between IPSL-CM5A-MR and MIROC-ESM-CHEM outputs under RCP4.5 for year 2070. The highest correlation coefficient value (*r* = 0.97) was given by HadGEM2-ES and NCAR-CCSM4 outputs under RCP2.6 for year 2070.

*S. campylacanthum* species binary outputs recorded the least correlation values (min *r* = 0.53) for both HadGEM2-ES and IPSL-CM5A-MR, and HadGEM2-ES and MIROC-ESM-CHEM outputs under RCP2.6 for year 2050. Maximum *r* value (*r* = 0.93) was given by IPSL-CM5A-MR and NCAR-CCSM4 outputs under RCP8.5 for year 2070. Binary outputs from HadGEM2-ES compared to those obtained from the rest of GCM were consistently below threshold *r* values (*r* < 0.70) under RCP2.6 for years 2050 and 2070, under RCP4.5 for year 2050 and under RCP8.5 for year 2070. We therefore concluded that HadGEM2-ES model prediction skill for *S. campylacanthum* species climate space was not satisfactory as compared to the rest of the GCMs (Fig 1e in S3 Appendix and Fig 1e in S4 Appendix) and hence was dropped from ensemble species distribution modelling of *S. campylacanthum* species. The six GCM model data were used for ensemble distribution modelling of rest of the species.

#### Ensemble species distribution changes

Ensemble species distribution changes showed an increase in habitat suitability for *L. camara* and *S. campylacanthum* species and a decline in habitat suitability for *C. decapetala, O. stricta, and S. didymobotrya* species for both 2050 and 2070 projection periods (Table 4; Figs 4, 5 and 6). *S. campylacanthum* species had the largest range expansion of ~33% of the total study area under RCP2.6 and RCP4.5 climate change scenario for both future periods 2050 and 2070. Its range expansion indicated a reduction of ~25% and ~30% under RCP8.5 for 2050 and 2070 respectively. *L. camara* had the second largest range expansion of ~12% under RCP2.6 and RCP4.5 for years 2050 and 2070. Its range expansion under RCP8.5 was ~14% and ~11% for years 2050 and 2070 respectively. *O. stricta* species had low range expansion of <1% under RCPs 2.6 and 4.5 for both future periods. Its lowest range expansion (~0.31%) was under RCP8.5 for the year 2050. Range expansion for *C. decapetala* on the other hand was ~1% for both future periods under all RCPs except for RCP8.5 for the year 2050 where the range expansion was the lowest (0.26%). *S. didymobotrya* species on the other hand recorded a range expansion of ~2% under all RCPs for both future periods. Its range expansion was towards the Aberdares national park and in Mukurwe-ini sub-county where initially absent in current distribution for both years (Figs 3 and 4).

**Fig 4.**
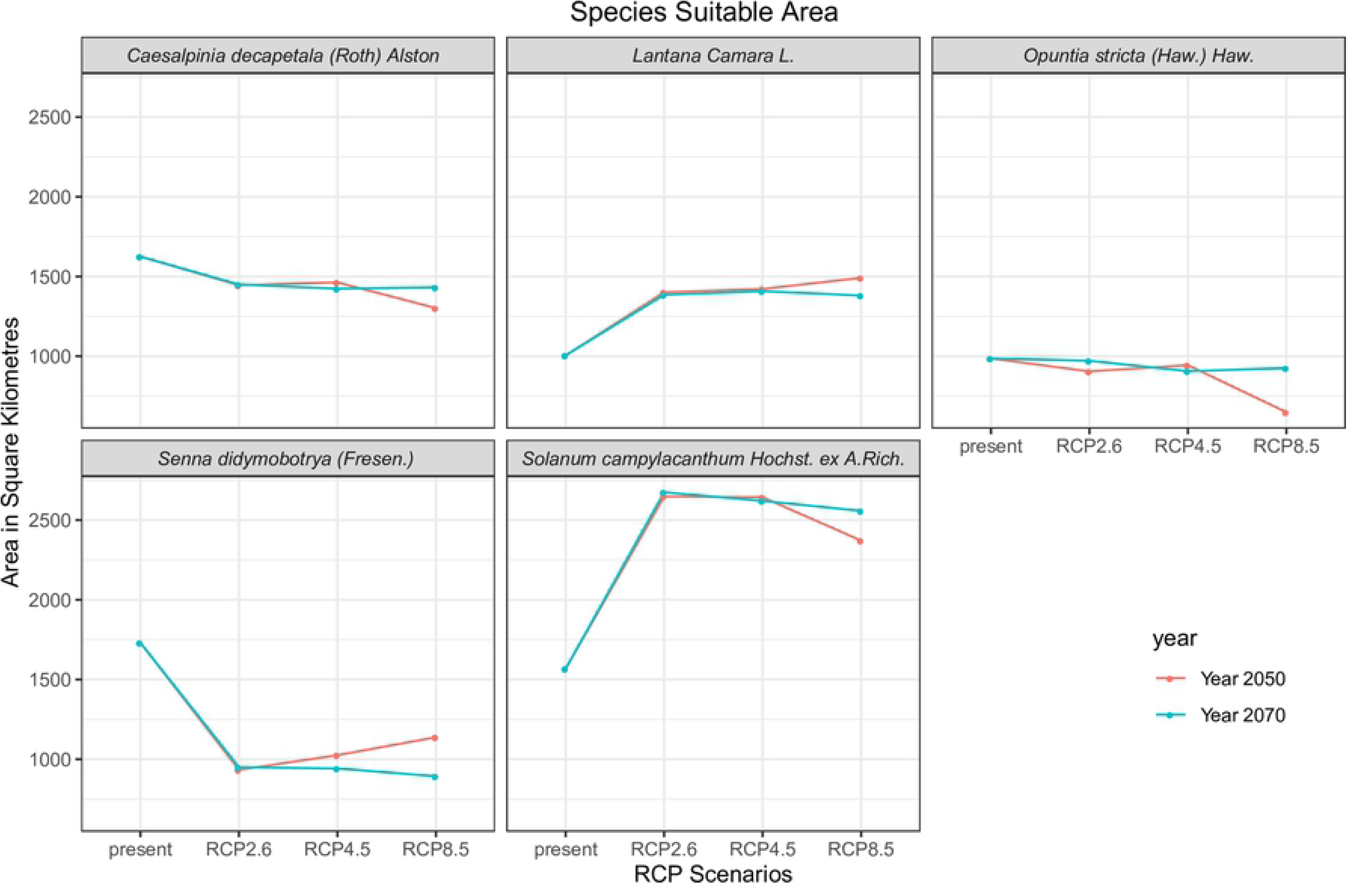
Predicted suitable areas under changing climate scenarios in square kilometres for the 5 study species.

**Fig 5.**
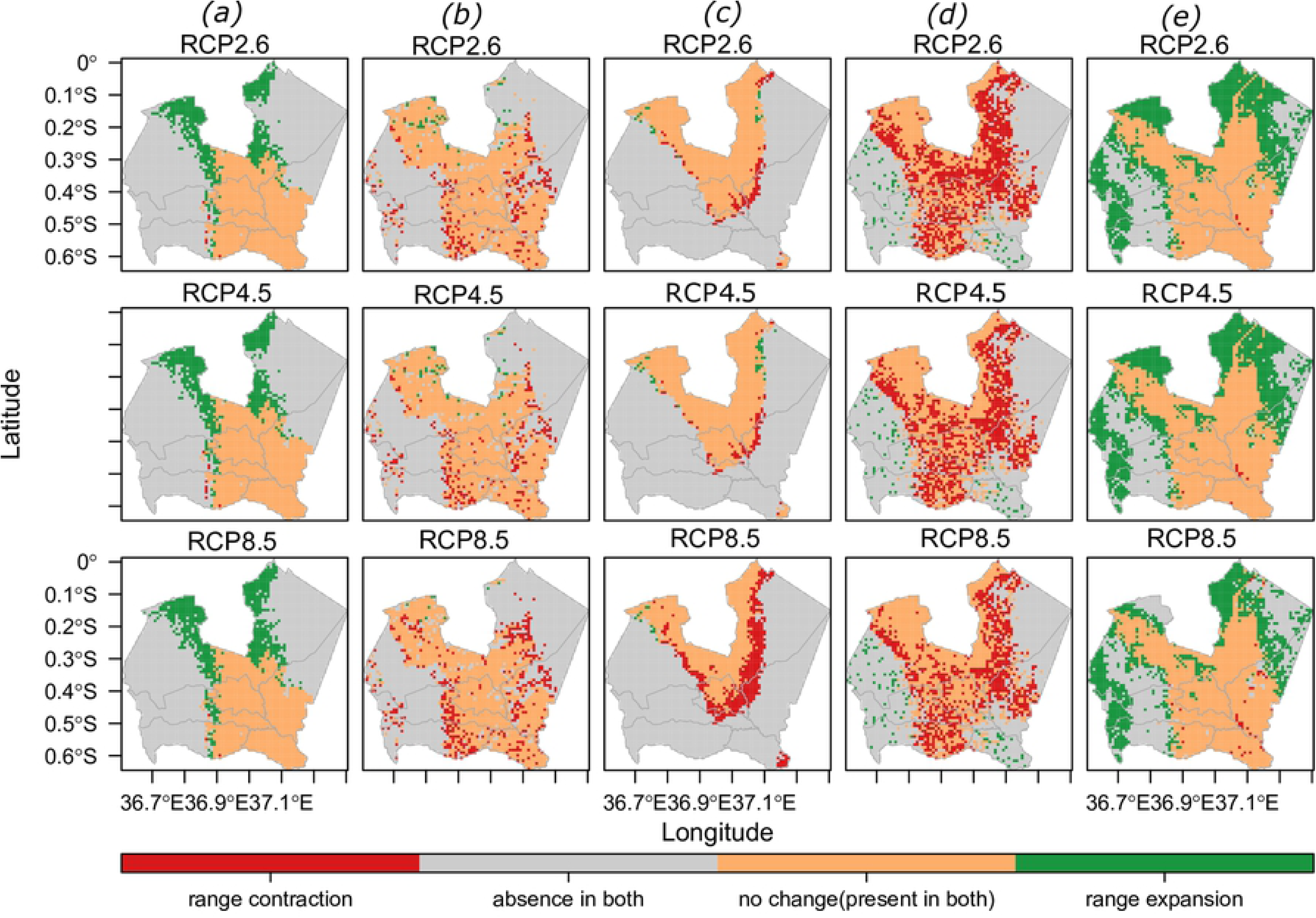
Ensemble species distribution changes between current and future climate scenarios under RCP2.6, RCP4.5, and RCP8.5 for the year 2050. (a) L. camara; (b) C. decapetala (Roth) Alston; (c) O. stricta; (d) S. didymobotrya; (e) S. campylacanthum Hochst. ex A. Rich.

**Fig 6.**
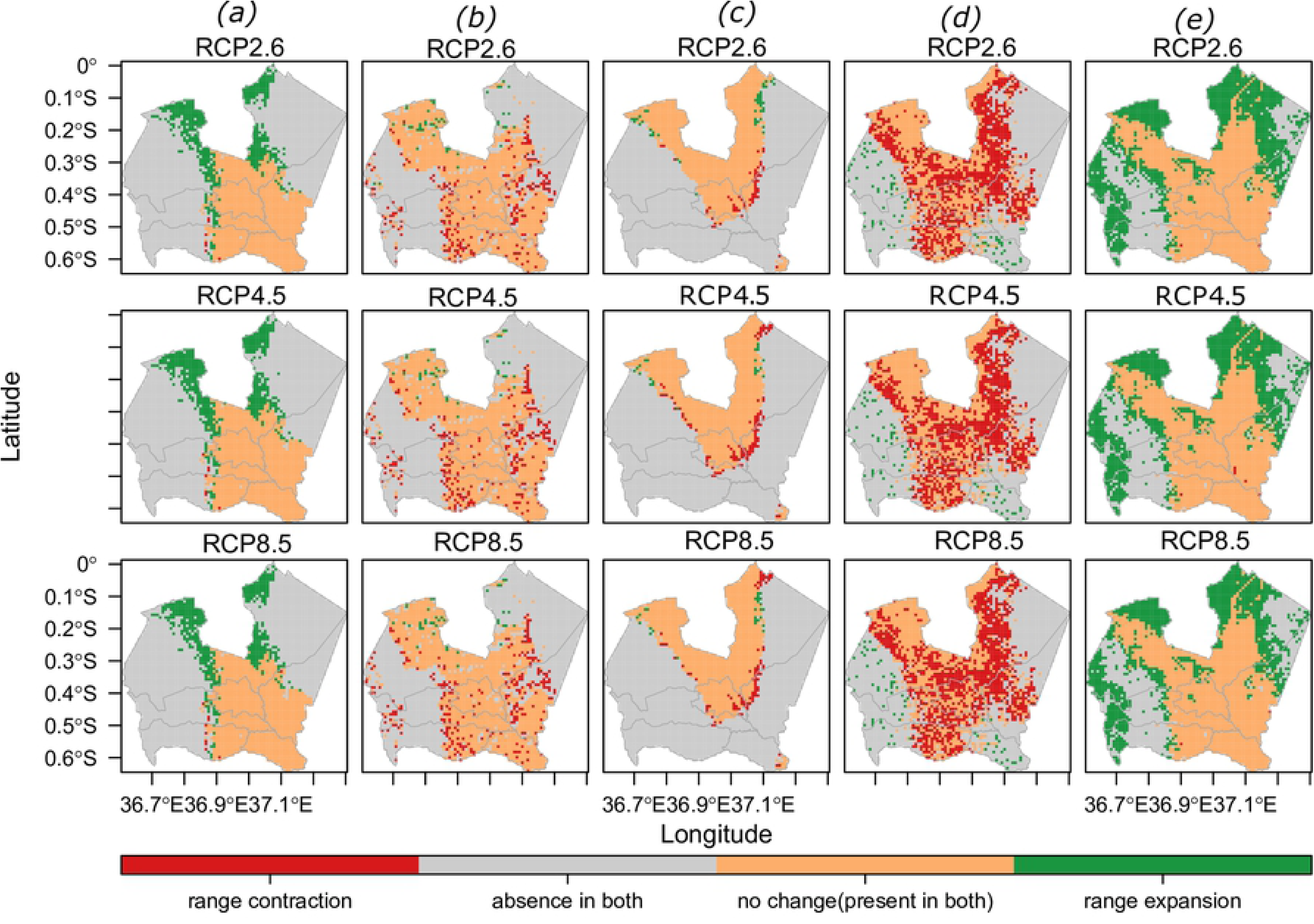
Ensemble species distribution changes between current and future climate scenarios under RCP2.6, RCP4.5, and RCP8.5 for the year 2070. (a) L. camara; (b) C. decapetala; (c) O. stricta; (d) S. didymobotrya; (e) S. campylacanthum

Although *S. didymobotrya* species had the largest suitability area (53%) among study species, its future habitat will contract by approximately half of its current habitat i.e. ~26% under RCP2.6 for both future periods. Its lowest range contraction (~20%) was under RCP8.5 for year 2050 and the largest (~27%) under RCP8.5 for year 2070. *C. decapetala* had the second largest range contraction after *S. didymobotrya* at ~6% under all RCPs and both future periods except under RCP8.5 for year 2050 which was at ~10%.

*O. stricta* distribution range had the highest range contraction at ~10% under RCP8.5 for year 2050 and the least at ~2% under RCP2.6 for year 2070. On the other hand, *L. camara* and *S. campylacanthum* had the lowest range contraction among the study species. The highest for *S. campylacanthum* was at ~1% under RCP8.5 for year2050 and the rest at <1%. *L. camara* range contraction was very low at ~0.03% – 0.08% under all RCPs for years 2050 and 2070.

Analysis of the species range shift indicated that *L. camara* species had the largest range shift (~8 – 10km) followed by *S. campylacanthum* (~6 – 7km). *S. didymobotrya’s* range shift distance was ~3 – 4km, *C. decapetala* ~1km while *O. stricta* had the largest range shift of ~5 km under RCP8.5 for the year 2050 and ~1km under the rest of RCPs for both years. All study species showed a North West range shift direction. Based on these results, it is evident that changing climatic conditions had little impact on *L. camara*, *S. campylacanthum* and *O. stricta species* suitable habitats within the study area. On the other hand, *S. didymobotrya* species suitable habitat was greatly reduced under changing climatic conditions.

## Discussion

Previous research has shown that climate change cause changes to the current environmental parameters of a given area [72] which in turn causes expansion of invasive species suitable areas [47]. Our study shows that *L. camara* and *S. campylacanthum* current distribution will persist and expand significantly to new habitats. Other than climate change enhancing range expansion, species may be subjected to an extinction trajectory. In our case, *S. didymobotrya* and *C. decapetala* have the largest range contraction among the study species. Despite *S. didymobotrya* being a native species and presumably well adapted to its environment, the impacts of changing climatic conditions will exacerbate its decline and a possible shift to new environments outside our study area. On the contrary, distribution of alien invasive species such as *L. camara*, *C. decapetala*, and *O. stricta* will persist under these climatic changes. This confirms that invasive species adapts well in diverse habitats with varying climatic conditions [8]. Although SDM outputs may give an under-prediction of a given study species due to difficulties in predicting species ability to evolve and adapt under changing conditions (Sinclair, White, and Newell 2010), our results confirms the role of climate change in enhancing bio-invasions in local natural environments [73].

Although climatic changing conditions will play a major role in reducing species habitats in future, continued decline and complete extinction is more likely to be influenced by habitat fragmentation over a longer period of time [74]. Land-use changes (habitat fragmentations) influences species biological processes as much as climate change does [35,72]. On the other hand, implying native species habitat decline as due to co-occurring alien invasive species may not suffice Gurevitch and Padilla [75] unless a more practical framework such as a six-threshold framework outlined in the work of Downey and Richardson [74] is used to assess the role of alien plants on native plants extinction. Contrary to perceived negative impacts in terms of habitat transformations by alien and native species Witt et al. [6], *S. didymobotrya* species has been found useful in African traditional medicine. For instance, Jeruto et al. [76] found that its stem and root extracts had high efficiencies in inhibiting fungus growth while Jeruto et al. [77] found that root bark extracts possess phytochemical properties that inhibit bacterial pathogens growth. In a conservation point of view, making policies that aim at conserving *S. didymobotrya* species especially in the wild would sustain availability of materials for development of alternative medicine. As such more studies on the species current and future habitats involving multiple county levels should be prioritized. As far as the other study species are concerned, their negative impacts on natural habitats may outweigh their positive benefits. Many studies have advocated for urgent control measures for species such as *L. camara* due to its negative impacts on socio-economic livelihoods and biodiversity [2,18]. Shackleton et al. [21] suggested urgent intervention measures on *O. stricta* species in Laikipia county due its impacts on annual economic losses per household among other negative impacts on rangelands. We foresee a similar scenario within our study area where *O. stricta* suitability in Kieni sub-county rangelands will affect pastoralism and wildlife conservation.

*S. campylacanthum* species, a native plant species, shows good adaptability to warming climatic conditions. Its gain in habitat includes parts of Mt. Kenya national park and forest reserve and the Aberdare national park and forest reserve (Figs 5 and 6). Extrapolation of climate envelope [72] to the entire study area shows new species habitat areas within unsampled protected areas (Fig 1) hence proving the usefulness of SDMs. Threats posed by alien plants invasion in high conservation areas is usually significant [74]. Our species distribution maps confirm possible proliferation of *C. decapetala*, *S. campylacanthum*, and *L*. *camara* species within protected areas both in current and future climates. These species had been cited as a nuisance in these protected areas by Kenya Forest Service [30]. Fragmented pixels within the Aberdare national park are predicted as suitable for *C. decapetala* species with a possible contraction in future climates (Figs 3 and 4). *L. camara* proliferation, as observed during field data collection, is pronounced within isolated forest conservation areas such as the Muringato nursery, Nyeri municipality and Nyeri forest conservation areas all of which fall within Nyeri Town sub-county (Fig 3b). Interests are high on *S. campylacanthum* species studies especially in conservation areas. For instance, a five-year research study by Pringle et al. [78] focusing on the effects of different sized mammalian herbivores including elephants, impala and dik-dik on *S. campylacanthum* species population within Mpala Research Centre in central Kenya indicated a complementary effect on the species population. While dik-dik reduced much of *S. campylacanthum* foliage, the impala and elephants contributed to seed dispersal hence complementing each other on sustaining species population. Since our study shows climate change will increase habitat range for *S. campylacanthum* species in conservation areas, its spread into new habitats will be fast due to availability of seed dispersers. Referring to the case of *S. campylacanthum* species, we have seen that while SDMs may provide an understanding to the distribution dynamics of invasive species, studies on other drivers of species population distribution are equally important.

For a local species SDM study e.g. areas of relatively small geographical extents, small-scale habitat attributes such as fine scale topography and vegetation metric such as NDMI and NDVI [79] are important. Such attributes are usually overlooked despite their profound importance [72]. Moreover, species distributions depend on additional biotic factors (e.g. ability of species to compete for nutrients) and dispersal factors of a given species [80] which were lacking in our SDM models. Nevertheless, most researchers use the readily available standard bioclimatic variables (mostly temperature and precipitation) as the only predictor variables for calibrating models for species future predictions e.g. Ashraf et al. [81]. Despite using predictor variables with 1km grain size for a small scale study [49], our current and future species distribution estimates provides information needed to activate management and monitoring actions in threatened habitats.

Fewer bioclimatic variables were selected as important (Table 2) a situation attributed to the fact that our study area is relatively small and possibly having little climate variations than that of a large spatial scale [49]. Topographic variables played a big role in species response and therefore cannot be ignored in small scale SDM studies. For instance, DEM variable shows decreasing low probability of occurrence for *L. camara* in higher elevated areas >1750m. Thapa et al. [35] also found *L. camara* range contracting in upper elevations and expanding in lower elevations in the Kailash Sacred Landscape, Nepal.

In conservation planning, one of the goals is to establish biogeographical patterns of a given species often through SDMs. Such efforts enable identification of species invasiveness, sites that need prioritization for rehabilitation as well as re-introduction of threatened species [72]. We have provided baseline information on distribution of study species within Nyeri county, a major milestone in moving towards maintaining healthy natural habitats and preventing climate change vulnerabilities such as wild fires, reduction of survival of endemic species and adverse effects on agricultural systems and water catchment areas brought about by uncontrolled spread of invasive species [82].

## Conclusions

Our study has demonstrated the importance of carrying out SDM at a local natural environment to establish where and which invasive species will lose, gain suitable habitats and persist with little change in future climate change scenarios. We have also shown that invasive species management programs in local natural environments ought to consider climate change aspects in addition to other topographic factors. Immediate actions are needed to avert possible losses of biodiversity due to persistence and future expansion of *L. camara* and *S. campylacanthum* invasive species within Nyeri county more so in the biodiversity hotspot areas. Additional work is needed to support local invasive species eradication programs more so on determination of their actual location and absolute quantification of area of occupancy. Moving forward, we intend to improve on this work by developing a mapping framework that utilizes species unique spectral indices to enhance rapid estimation of fractional cover maps as well as derivation of essential biodiversity variables from remote sensing imageries for building accurate SDMs at local scale levels.

## Acknowledgements

The study was part of PhD research by Julius M Waititu. We would like to covey our gratitude to the Kenya Forest Service for permitting research within Nyeri forests ecosystem. We would also want to thank Mr. Peter Kariuki, Manager Muringato Nyeri conservation area for his assistance in accessing Nyeri forest stations. The other gratitude goes to Mr. L. Kortom, Ms. A. Theuri and Mr. J. Njeru for their assistance in field data collection.

## Author Contributions

Conceived and conceptualized the idea: JMW, CNM, AWS.

Collected data, performed preliminary analysis and interpretation of results and wrote the first draft of the manuscript: JMW.

Reviewed and gave corrections to the manuscript and improved the ideas in it through discussions. CNM and AWS.

## Supporting information

**S1 Appendix.** S**ampled grid cells, description of model predictor variables, minimum SDM records and model evaluation metrics.** (PDF)

**S2 Appendix. Predictor Variable Relative Importance and Species Response Curves.**(PDF)

**S3 Appendix. Species distribution change maps between current and future climate scenarios for the year 2050.** (PDF)

**S4 Appendix. Species distribution change maps between current and future climate scenarios for the year 2070.** (PDF).

**S5 Table. Pearson correlation coefficient (r) and significant differences (p < .001) for GCM models future distribution maps output.** (PDF)

**S6 Dataset. Rarefied field species presence records.** (ZIP)

## References

1. Richardson DM, Pysek P, Rejmanek M, Barbour MG, Panetta FD, West CJ. Naturalization and invasion of alien plants: concepts and definitions. Divers Distrib [Internet]. 2000 Mar;6(2):93–107. Available from: http://doi.wiley.com/10.1046/j.1472-4642.2000.00083.x

2. Subhashni T, Lalit K. Impacts of climate change on invasive Lantana camara L. distribution in South Africa. African J Environ Sci Technol [Internet]. 2014;8(6):391–400. Available from: http://academicjournals.org/journal/AJEST/article-abstract/D9EF7A045968

3. United Nations. United Nations, Transforming Our World: The 2030 Agenda for Sustainable Development [Internet]. 2015 [cited 2018 Oct 8]. Available from:https://sustainabledevelopment.un.org/content/documents/21252030 Agenda for Sustainable Development web.pdf

4. Radosevich SR, Stubbs MM, Ghersa CM. Plant invasions-process and patterns. Weed Sci. 2003;51:254–9.

5. Royimani L, Mutanga O, Odindi J, Dube T, Matongera TN. Advancements in satellite remote sensing for mapping and monitoring of alien invasive plant species (AIPs). Phys Chem Earth, Parts A/B/C [Internet]. 2018 Dec [cited 2019 Jan 25]; Available from:https://linkinghub.elsevier.com/retrieve/pii/S1474706518301128

6. Witt A, Beale T, van Wilgen BW. An assessment of the distribution and potential ecological impacts of invasive alien plant species in eastern Africa. Trans R Soc South Africa [Internet]. 2018 Sep 2;73(3):217–36. Available from:https://www.tandfonline.com/doi/full/10.1080/0035919X.2018.1529003

7. Richardson DM, Pyšek P. What is an Invasive Species? [Internet]. Crop Protection Compendium. 2004 [cited 2020 Mar 6]. p. 17. Available from:https://www.cabi.org/isc/FullTextPDF/2009/20093238299.pdf

8. Hellmann JJ, Byers JE, Bierwagen BG, Dukes JS. Five potential consequences of climate change for invasive species. Conserv Biol. 2008;22(3):534–43.

9. IPCC. Climate Change 2014: Synthesis Report. Contribution of Working Groups I, II and III to the Fifth Assessment Report of the Intergovernmental Panel on Climate Change [Core Writing Team, R.K. Pachauri and L.A. Meyer (eds.)] [Internet]. 2014. Available from: ipcc.ch/site/assets/uploads/2018/05/SYR_AR5_FINAL_full_wcover.pdf

10. Moss RH, Edmonds JA, Hibbard KA, Manning MR, Rose SK, van Vuuren DP, et al. The next generation of scenarios for climate change research and assessment. Nature [Internet]. 2010 Feb;463(7282):747–56. Available from: http://dx.doi.org/10.1038/nature08823

11. IPCC, Allen M, Babiker M, Chen Y, de Coninck H, Connors S, et al. Summary for Policymakers. In: Global warming of 1.5°C. An IPCC Special Report. In 2018.

12. Hejda M, Pyšek P, Jarošík V. Impact of invasive plants on the species richness, diversity and composition of invaded communities. J Ecol. 2009;97(3):393–403.

13. van Proosdij ASJ, Sosef MSM, Wieringa JJ, Raes N. Minimum required number of specimen records to develop accurate species distribution models. Ecography (Cop) [Internet]. 2016 Jun;39(6):542–52. Available from: http://doi.wiley.com/10.1111/ecog.01509

14. Rivera ÓR de, López-Quílez A. Development and Comparison of Species Distribution Models for Forest Inventories. ISPRS Int J Geo-Information [Internet]. 2017 Jun 16 [cited 2018 Dec 20];6(6):176. Available from: http://www.mdpi.com/2220-9964/6/6/176

15. Shabani F, Kumar L, Ahmadi M. A comparison of absolute performance of different correlative and mechanistic species distribution models in an independent area. Ecol Evol. 2016;6(16):5973–86.

16. Taylor S, Kumar L. Sensitivity Analysis of CLIMEX Parameters in Modelling Potential Distribution of Lantana camara L. PLoS One [Internet]. 2012 [cited 2018 Oct 8];7(7):40969. Available from: www.plosone.org

17. Truong TTA, Hardy GESJ, Andrew ME. Contemporary Remotely Sensed Data Products Refine Invasive Plants Risk Mapping in Data Poor Regions. Front Plant Sci [Internet]. 2017;8(May). Available from: http://journal.frontiersin.org/article/10.3389/fpls.2017.00770/full

18. Shackleton RT, Witt ABR, Aool W, Pratt CF. Distribution of the invasive alien weed, Lantana camara, and its ecological and livelihood impacts in eastern Africa. African J Range Forage Sci. 2017;34(1):1–11.

19. Brummer TJ, Maxwell BD, Higgs MD, Rew LJ. Implementing and interpreting local-scale invasive species distribution models. Franklin J, editor. Divers Distrib [Internet]. 2013 Aug;19(8):919–32. Available from: http://doi.wiley.com/10.1111/ddi.12043

20. Lowe S, Browne M, Boudjelas S, Poorter M De. 100 of the World’s Worst Invasive Alien Species A selection from the Global Invasive Species Database. [Internet]. 2000. Available from: www.issg.org/booklet.pdf

21. Shackleton RT, Witt ABR, Piroris FM, van Wilgen BW. Distribution and socio-ecological impacts of the invasive alien cactus Opuntia stricta in eastern Africa. Biol Invasions. 2017;19(8):2427–41.

22. Gichua M, Njoroge G, Shitanda D, Ward D. Invasive species in east africa: current status for informed policy decisions and management. JAGST [Internet]. 2013 [cited 2018 Oct 8];15(1):45–55. Available from: http://journals.jkuat.ac.ke/index.php/jagst/article/viewFile/1015/824

23. Seburanga JL. Black Wattle (Acacia mearnsii De Wild.) in Rwanda’s Forestry: Implications for Nature Conservation. J Sustain For. 2015;34(3):276–99.

24. Byrne MJ, Witkowski ETF, Kalibbala FN. A Review of Recent Efforts at Biological Control of Caesalpinia decapetala (Roth) Alston (Fabaceae) in South Africa. African Entomol. 2011;19(2):247–57.

25. Witt A, Luke Q. Guide to the naturalized and invasive plants of Eastern Africa [Internet]. Witt A, Luke Q, editors. Wallingford, UK: CABI; 2017. vi + 601 pp. Available from: http://www.cabi.org/cabebooks/ebook/20173158959

26. Government of the Republic of Kenya. Second Medium Term Plan, 2013 – 2017 [Internet]. Nairobi; 2013 [cited 2018 Oct 17]. Available from: http://vision2030.go.ke/inc/uploads/2018/06/Second-Medium-Term-Plan-2013-2017.pdf

27. UNDP and County Government of Marsabit. Revised first county integrated development plan [Internet]. 2013. Available from: http://www.ke.undp.org/content/dam/kenya/docs/Democratic Governance/Marsabit County Revised CIDP.pdf

28. Government of Kenya. Nyeri County Intergrated Development Plan 2018-2022 [Internet]. 2018. Available from: http://www.nyeri.go.ke/wp-content/uploads/2017/01/County-Govt-of-Nyeri-CIDP.pdf

29. Gachathi F, Ngugi J, Omondi S. Useful trees suitable for central highlands eco-region [Internet]. Central Highlands Eco-region Research Programme, Kenya Forestry Research Institute (KEFRI); 2014 [cited 2020 Feb 3]. Available from:https://www.kefri.org/PDF/Leaflets/USEFULTREESSUITABLEFORCENTRALHIGHLANDSECO-REGION.pdf

30. Kenya Forest Service. Aberdare forest reserve managment plan [Internet]. 2010 [cited 2020 Mar 4]. p. 94. Available from: http://www.kenyaforestservice.org/documents/Aberdare.pdf

31. Shuster WD, Herms CP, Frey MN, Doohan DJ, Cardina J. Comparison of survey methods for an invasive plant at the subwatershed level. Biol Invasions. 2005;7(3):393–403.

32. Meunier G, Lavoie C. Roads as Corridors for Invasive Plant Species: New Evidence from Smooth Bedstraw (Galium mollugo). Invasive Plant Sci Manag. 2012;5(1):92–100.

33. Von Der Lippe M, Kowarik I. Long-distance dispersal of plants by vehicles as a driver of plant invasions. Conserv Biol. 2007;21(4):986–96.

34. Dillon WW, Lieurance D, Hiatt DT, Clay K, Flory SL. Native and invasive woody species differentially respond to forest edges and forest successional age. Forests. 2018;9(7):1–17.

35. Thapa S, Chitale V, Rijal SJ, Bisht N, Shrestha BB. Understanding the dynamics in distribution of invasive alien plant species under predicted climate change in Western Himalaya. Liu J, editor. PLoS One [Internet]. 2018 Apr 17;13(4):e0195752. Available from:https://dx.plos.org/10.1371/journal.pone.0195752

36. Henderson L. Invasive, naturalized and casual alien plants in southern Africa: a sum-mary based on the Southern African Plant Invaders Atlas (SAPIA). Bothalia [Internet]. 2007;37(2):215–48. Available from: http://abcjournal.org/index.php/ABC/article/view/322

37. Wabuyele E, Lusweti A, Bisikwa J, Kyenune G, Clark K, Lotter WD, et al. A Roadside Survey of the Invasive Weed Parthenium hysterophorus (Asteraceae) in East Africa. J East African Nat Hist [Internet]. 2014;103(1):49–57. Available from: http://www.bioone.org/doi/10.2982/028.103.0105

38. McPherson JM, Jetz W, Rogers DJ. The effects of species’ range sizes on the accuracy of distribution models: ecological phenomenon or statistical artefact? J Appl Ecol [Internet]. 2004 Sep 30;41(5):811–23. Available from: http://doi.wiley.com/10.1111/j.0021-8901.2004.00943.x

39. Merow C, Smith MJ, Silander JA. A practical guide to MaxEnt for modeling species’ distributions: What it does, and why inputs and settings matter. Ecography (Cop). 2013;36(10):1058–69.

40. Fick SE, Hijmans RJ. Worldclim 2: New 1-km spatial resolution climate surfaces for global land areas. [Internet]. International Journal of Climatology. 2017 [cited 2019 Feb 2]. Available from: http://worldclim.org/version2

41. Trabucco A, Zomer RJ. Global Aridity Index and Potential Evapo-Transpiration (ET0) Climate Database v2. CGIAR Consortium for Spatial Information (CGIAR-CSI). [Internet]. Published Online. 2018 [cited 2020 Mar 2]. Available from:https://doi.org/10.6084/m9.figshare.7504448.v3

42. Dataset ASF DAAC. ALOS PALSAR_Radiometric_Terrain_Corrected_high_res; Includes Material ©JAXA/METI [2007]. 2007.

43. ESA. Land Cover CCI Product User Guide Version 2. Tech. Rep. [Internet]. 2017 [cited 2020 Feb 3]. Available from: http://maps.elie.ucl.ac.be/CCI/viewer/download/ESACCI-LC-Ph2-PUGv2_2.0.pdf

44. Hengl T, Mendes de Jesus J, Heuvelink GBM, Ruiperez Gonzalez M, Kilibarda M, Blagotić A, et al. SoilGrids250m: Global gridded soil information based on machine learning. Bond-Lamberty B, editor. PLoS One [Internet]. 2017 Feb 16;12(2):e0169748. Available from:https://dx.plos.org/10.1371/journal.pone.0169748

45. Navarro-Racines C, Tarapues J, Thornton P, Jarvis A, Ramirez-Villegas J. High-resolution and bias-corrected CMIP5 projections for climate change impact assessments. Sci Data [Internet]. 2020 Dec 20;7(1):7. Available from: http://www.nature.com/articles/s41597-019-0343-8

46. McSweeney CF, Jones RG, Lee RW, Rowell DP. Selecting CMIP5 GCMs for downscaling over multiple regions. Clim Dyn. 2015;

47. Shrestha UB, Shrestha B B. Climate change amplifies plant invasion hotspots in Nepal. Vaclavik T, editor. Divers Distrib [Internet]. 2019 Oct 2;25(10):1599–612. Available from:https://onlinelibrary.wiley.com/doi/abs/10.1111/ddi.12963

48. Aguirre-Gutiérrez J, van Treuren R, Hoekstra R, van Hintum TJL. Crop wild relatives range shifts and conservation in Europe under climate change. Divers Distrib. 2017;23(7):739–50.

49. Manzoor SA, Griffiths G, Lukac M. Species distribution model transferability and model grain size - finer may not always be better. Sci Rep [Internet]. 2018;8(1):1–9. Available from: http://dx.doi.org/10.1038/s41598-018-25437-1

50. Jackson LS, Carslaw N, Carslaw DC, Emmerson KM. Modelling trends in OH radical concentrations using generalized additive models. Atmos Chem Phys. 2009;9(6):2021–33.

51. R Core Team. R: A Language and Environment for Statistical Computing [Internet]. Vienna, Austria: R Foundation for Statistical Computing; 2013. Available from: http://www.r-project.org/

52. Naimi B. Package “usdm”. Uncertainty Analysis for Species Distribution Models. R-Cran. 2017;

53. Naimi B, Araújo MB. sdm: a reproducible and extensible R platform for species distribution modelling. Ecography (Cop) [Internet]. 2016 Apr;39(4):368–75. Available from: http://doi.wiley.com/10.1111/ecog.01881

54. Brown JL. SDMtoolbox: A python-based GIS toolkit for landscape genetic, biogeographic and species distribution model analyses. Methods Ecol Evol. 2014;5(7):694–700.

55. Elith J, Kearney M, Phillips S. The art of modelling range-shifting species. Methods Ecol Evol. 2010;1(4):330–42.

56. Barbet-Massin M, Jiguet F, Albert CH, Thuiller W. Selecting pseudo-absences for species distribution models: How, where and how many? Methods Ecol Evol. 2012;3(2):327–38.

57. Liu C, Newell G, White M. On the selection of thresholds for predicting species occurrence with presence-only data. Ecol Evol. 2016;6(1):337–48.

58. Phillips SJ, Anderson RP, Schapire RE. Maximum entropy modeling of species geographic distributions. Ecol Modell [Internet]. 2006 [cited 2019 Feb 2];190:231–59. Available from: http://webpages.icav.up.pt/ptdc/BIA-BIC/110587/2009/Papers/20.pdf

59. Breiman L. Random Forests. Mach Learn [Internet]. 2001;45(1):5–32. Available from: http://link.springer.com/10.1023/A:1010933404324

60. Zhang L, Huettmann F, Liu S, Sun P, Yu Z, Zhang X, et al. Classification and regression with random forests as a standard method for presence-only data SDMs: A future conservation example using China tree species. Ecol Inform [Internet]. 2019;52(1):46–56. Available from:https://doi.org/10.1016/j.ecoinf.2019.05.003

61. Shabani F, Kumar L, Ahmadi M. A comparison of absolute performance of different correlative and mechanistic species distribution models in an independent area. Ecol Evol [Internet]. 2016 Aug [cited 2018 Dec 20];6(16):5973–86. Available from: http://doi.wiley.com/10.1002/ece3.2332

62. Radosavljevic A, Anderson RP. Making better Maxent models of species distributions:Complexity, overfitting and evaluation. J Biogeogr. 2014;41(4):629–43.

63. Liaw A, Wiener M. Classification and Regression by randomForest. R News. 2002;

64. Phillips SJ.A Brief Tutorial on Maxent [Internet]. 2017 [cited 2020 Dec 3]. Available from:https://biodiversityinformatics.amnh.org/open_source/maxent/

65. Pearce J, Ferrier S. Evaluating the predictive performance of habitat models developed using logistic regression. Ecol Modell. 2000;133(3):225–45.

66. Allouche O, Tsoar A, Kadmon R. Assessing the accuracy of species distribution models: Prevalence, kappa and the true skill statistic (TSS). J Appl Ecol. 2006;43(6):1223–32.

67. Liu C, White M, Newell G. Selecting thresholds for the prediction of species occurrence with presence-only data. J Biogeogr. 2013;40(4):778–89.

68. Raes N, ter Steege H. A null-model for significance testing of presence-only species distribution models. Ecography (Cop) [Internet]. 2007 Oct;30(5):727–36. Available from: http://doi.wiley.com/10.1111/j.2007.0906-7590.05041.x

69. Hijmans RJ, Phillips S, Leathwick J, Elith J. Package ‘dismo’’ - Species Distribution Modeling.’ CRAN Repository. 2017.

70. Harrell FE, Dupont C. Package ‘Hmisc’: Harrell Miscellaneous. R Top Doc. 2016;

71. Zhang L, Liu S, Sun P, Wang T, Wang G, Zhang X, et al. Consensus forecasting of species distributions: The effects of niche model performance and niche properties. PLoS One. 2015;10(3).

72. Sinclair SJ, White MD, Newell GR. How useful are species distribution models for managing biodiversity under future climates? Ecol Soc. 2010;15(1).

73. Shrestha UB, Sharma KP, Devkota A, Siwakoti M, Shrestha BB. Potential impact of climate change on the distribution of six invasive alien plants in Nepal. Ecol Indic. 2018;95:99–107.

74. Downey PO, Richardson DM. Alien plant invasions and native plant extinctions: a six-threshold framework. AoB Plants. 2016;8:plw047.

75. Gurevitch J, Padilla DK. Are invasive species a major cause of extinctions? Trends Ecol Evol. 2004;19(9):470–4.

76. Jeruto P, Arama PF, Anyango B, Akenga T, Nyunja R, Khasabuli D. In vitro antifungal activity of methanolic extracts of different senna didymobotrya (fresen.) H.S. Irwin & barneby plant parts. African J Tradit Complement Altern Med. 2016;

77. Jeruto P, Arama PF, Anyango B, Maroa G. Phytochemical screening and antibacterial investigations of crude methanol extracts of Senna didymobotrya (Fresen.) H. S. Irwin & Barneby. J Appl Biosci [Internet]. 2017 Sep 27;114(1):11357. Available from:https://www.ajol.info/index.php/jab/article/view/161557

78. Pringle RM, Goheen JR, Palmer TM, Charles GK, DeFranco E, Hohbein R, et al. Low functional redundancy among mammalian browsers in regulating an encroaching shrub (Solanum campylacanthum) in African savannah. Proc R Soc B Biol Sci. 2014;

79. Kearney M, Porter W. Mechanistic niche modelling: combining physiological and spatial data to predict species’ ranges. Ecol Lett [Internet]. 2009 Apr;12(4):334–50. Available from: http://doi.wiley.com/10.1111/j.1461-0248.2008.01277.x

80. Pearson RG. Species’ Distribution Modeling for Conservation Educators and Practitioners. Lessons Conserv [Internet]. 2007;3:54–89. Available from: http://ncep.amnh.org/linc

81. Ashraf U, Peterson AT, Chaudhry MN, Ashraf I, Saqib Z, Ahmad SR, et al. Ecological niche model comparison under different climate scenarios: A case study of Olea spp. in Asia. Ecosphere. 2017;8(5):1–13.

82. IUCN. Invasive alien species and climate change [Internet]. 2017 [cited 2019 Feb 2]. Available from:https://www.iucn.org/sites/dev/files/ias_and_climate_change_issues_brief_final.pdf

